# Assessing the causal role of body mass index on cardiovascular health in young adults: Mendelian randomization and recall-by-genotype analyses

**DOI:** 10.1101/112912

**Authors:** Kaitlin H. Wade, Scott T. Chiesa, Alun D. Hughes, Nish Chaturvedi, Marietta Charakida, Alicja Rapala, Vivek Muthurangu, Tauseef Khan, Nicholas Finer, Naveed Sattar, Laura D. Howe, Abigail Fraser, Debbie A. Lawlor, George Davey Smith, John E. Deanfield, Nicholas J. Timpson

## Abstract

**Background:** Mendelian randomization (MR) studies of body mass index (BMI) and cardiovascular health in mid-to-late life suggest causal relationships, but the nature of these has not been explored systematically at younger ages. Using complementary MR and recall-by-genotype (RbG) methodologies, our objective was to estimate the causal effect of BMI on detailed measures of cardiovascular health in a population of young healthy adults.

**Methods and Findings:** Data from the Avon Longitudinal Study of Parents and Children were used. For MR analyses, a genetic risk score (GRS) comprising 97 independent single nucleotide polymorphisms (SNPs) and constructed using external weighting was used as an instrument to test the causal effect of each unit increase in BMI (kg/m^2^) on selected cardiovascular phenotypes measured at age 17 (N=7909). An independent enriched sample from the same cohort participated in a RbG study at age 21, which enabled more detailed cardiovascular phenotyping (N=418; 191/227 from the lower/upper ∼30% of a genome-wide GRS distribution predicting variation in BMI). The causal effect of BMI on the additional cardiovascular phenotypes was assessed by comparing the two recalled groups. Difference in mean BMI between RbG groups was 3.85kg/m^2^ (95% CI: 2.53, 4.63; *P*=6.09×10^11^). In both MR and RbG analyses, results indicated that higher BMI causes higher blood pressure (BP) and left ventricular mass (indexed to height^2.7^, LVMI) in young adults (e.g. difference in LVMI per kg/m^2^ using MR: 1.07g/m^2.7^; 95% CI: 0.62, 1.52; P=3.87×10^−06^ and per 3.58kg/m^2^ using RbG: 1.65g/m^2.7^ 95% CI: 0.83, 2.47; P=0.0001). Additionally, RbG results indicated a causal role of higher BMI on higher stroke volume (SV; difference per 3.58kg/m^2^: 1.49ml/m^2.04^; 95% CI: 0.62, 2.35; *P*=0.001) and cardiac output (CO; difference per 3.58kg/m^2^: 0.11l /min/m^1.83^; 95% CI: 0.03, 0.19; *P*=0.01). Neither analysis supported a causal role of higher BMI on heart rate.

**Conclusions:** Complementary MR and RbG causal methodologies, together with a range of appropriate sensitivity analyses, showed that higher BMI is likely to cause worse cardiovascular health, specifically higher BP and LVMI, even in youth. These consistent results support efforts to prevent or reverse obesity in the young.

## Introduction

Higher body mass index (BMI) in adulthood is likely to be causally associated with numerous cardiovascular risk factors and disease outcomes[1-6]. Whilst chronic conditions may have origins early on, these relationships have been assessed predominantly from the 5^th^ decade of life[7]. It is assumed that these relationships reflect long-term exposure to adiposity and other co-morbidities, which result in adverse structural and functional cardiovascular changes that are different from adaptations encountered earlier in the disease’s evolution.

Numerous observational studies have reported associations between higher BMI and the presence of various subclinical markers of cardiovascular disease (CVD) even during young adulthood[8-11]. However, these observational study designs preclude a distinction between correlation and causation due to issues such as confounding or possible reverse causation (particularly in studies of older adults, whereby the ‘outcome’ is responsible for the variation in the ‘exposure’). Whilst recent results from patients who have had bariatric surgery provide supportive evidence for a causal role of greater adiposity on risk of major cardiovascular events[4, 5], no large studies have assessed the causal impact between BMI and cardiovascular phenotypes in early life where risk emerges.

One method for establishing evidence for causal relationships between an exposure and outcome of interest is Mendelian Randomization (MR), a technique which uses genetic variations as instrumental variables (IVs) in observational epidemiological studies[12, 13]. Due to the technique’s requirement for relatively large sample sizes to provide adequate statistical power, however, MR studies traditionally use either routinely collected clinical measures or data generated from high-throughput technologies. Detailed and precise subclinical measures of early structural and functional vascular adaptations are not commonly carried out in large population studies, as these are expensive, time-consuming and require highly skilled operators. This has limited their suitability for analyses using MR methodology.

Recall-by-genotype (RbG) studies are an innovative extension of MR designed to improve study efficiency through the creation of subgroups selected based on genotypes possessing known correlations with exposures of interest (e.g. BMI), rather than random sampling based on extremes of BMI itself.The added statistical power of this technique enables the efficient collection of extremely precise phenotypic data that may be otherwise impractical at the scale necessary for MR analyses[14, 15].

Using data from the Avon Longitudinal Study of Parents and Children (ALSPAC), we aimed to employ these two complementary analytical approaches (MR and RbG), alongside conventional multivariable regression analyses, to support the hypothesis that BMI causally influences variations in multiple clinically relevant measures of cardiovascular structure and function in adolescence and early adulthood, when risk emerges.

## METHODS

### Cohort description

The Avon Longitudinal Study of Parents and Children (ALSPAC) is a prospective birth cohort study investigating factors that influence normal childhood development and growth. The cohort and study design have been described in detail previously[16, 17] and are available at the ALSPAC website (http://www.alspac.bris.ac.uk). The study website contains details of all data that is available through a fully searchable data dictionary (http://www.bris.ac.uk/alspac/researchers/data-access/data-dictionary). Briefly, 14541 pregnant women resident in a defined area of the South West of England, with an expected delivery date of 1^st^ of April 1991 - 31^st^ of December 1992 were enrolled to the cohort. Of these, 13988 live-born children who were still alive 1 year later have been followed-up to date with regular questionnaires and clinical measures, providing behavioural, lifestyle and biological data. Ethical approval for the study was obtained from the ALSPAC Ethics and Law Committee and the Local Research Ethics Committee and written informed consent was obtained from both the parent/guardian and, after the age of 16, children provided written assent.

### Study design

The two complementary analytical approaches (MR and RbG) were used to support the hypothesis that BMI causally influences variations in multiple clinically relevant measures of cardiovascular structure and function in adolescence and early adulthood (Figure 1). Firstly, we used a genetic risk score (GRS) comprising 97 BMI-associated single nucleotide polymorphisms (SNPs) constructed using external weighting as an IV within an MR framework to investigate the causal effect of BMI on a range of vascular measures collected from 17-year-old participants[18].

**Figure 1.**
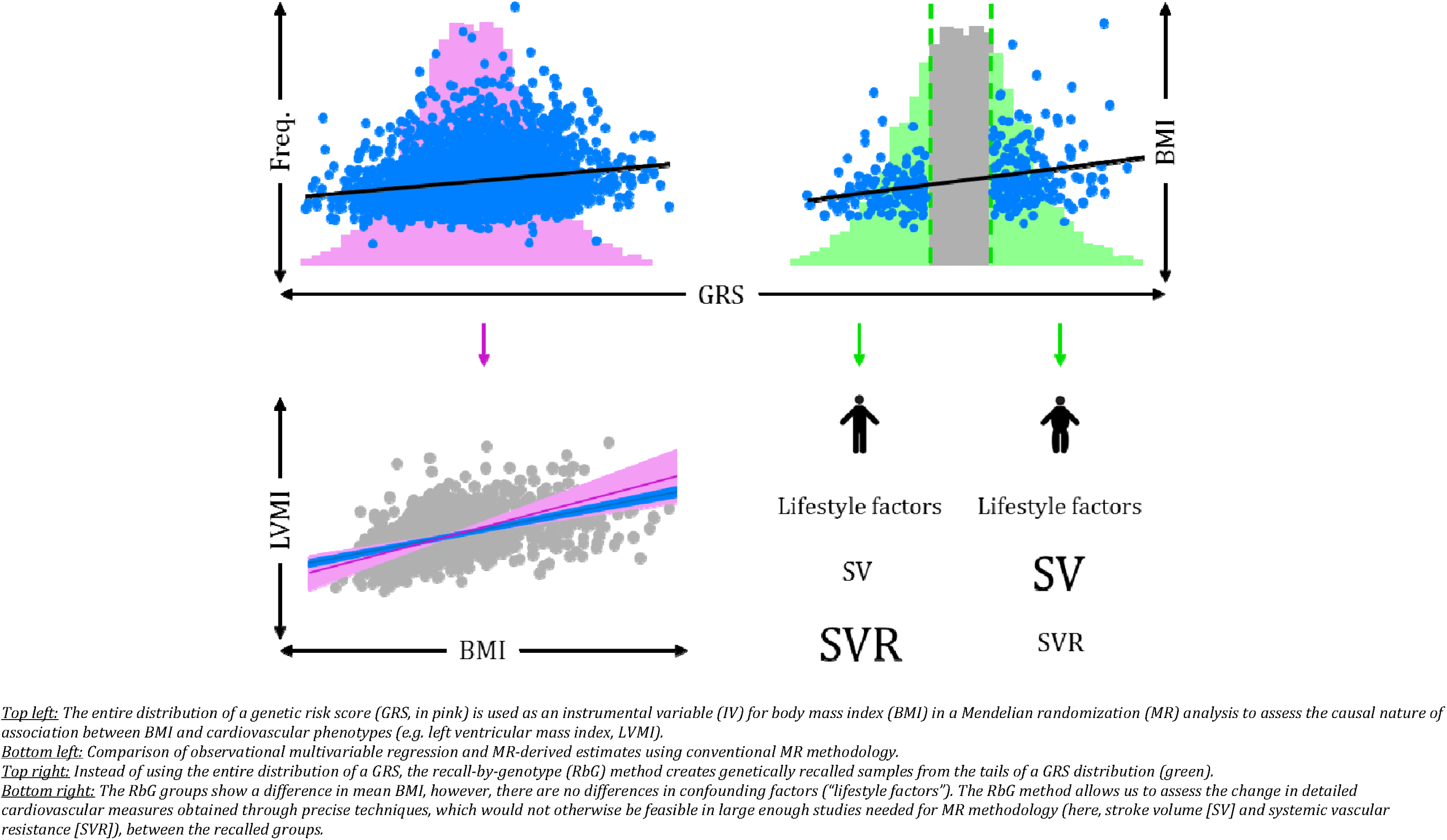
Mendelian randomization (left) and recall-by-genotype (right) methodologies.

In parallel with this approach, we examined the ability of a highly-powered RbG design to reproduce these findings and explore their underlying mechanisms further through the extensive phenotyping of a smaller group of independent individuals, recalled specifically on a genome-wide GRS distribution[19].

Of those with full genetic data and consent (N=8,350), individuals were invited to the RbG study based on the lower and upper ∼30% of a genome-wide GRS distribution, constructed from results from a genome-wide association study (GWAS) of BMI conducted by Speliotes *et al*.[19]. Individuals were recalled based on their appearance in the sampling groups, from the most extreme to the least to maximise power and difference in BMI. Of those invited (N=4602), 419 individuals were successfully recalled across the distribution of invited sampling groups (Supplementary Methods).

We excluded data of all females who were pregnant or individuals who had diabetes at both the 17-year clinic (N=7 pregnancies and 15 diabetics) and the 21-year recall (N=1 pregnancy and 0 diabetics). After exclusions, 7,909 individuals at age 17 were included in MR-analyses, which used a GRS comprising 97 SNPs (constructed using external weighting) shown to be associated with BMI from a large-scale GWAS[18]. The independent sample of 418 individuals were used in the 21-year RbG analyses (Figure 2).

**Figure 2.**
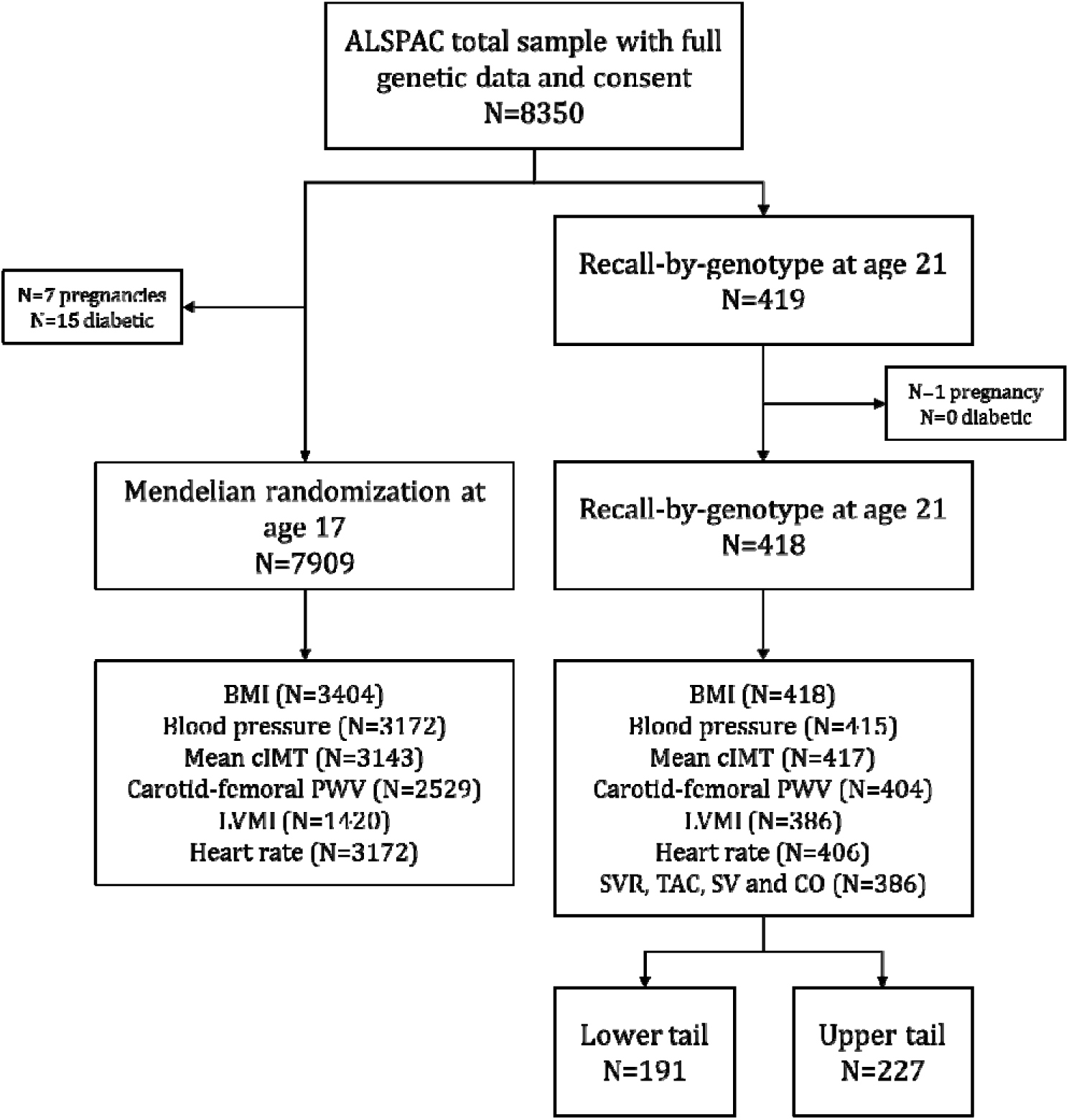
Flow of samples 153 used for Mendelian randomization and recall-by-genotype studies. *The total number of individuals in ALSPAC with full genetic data and consent was 8,350. Of these, 7,909 individuals were used in MR analyses at age 17, which used a GRS comprising 97 SNPs (and constructed using external weighting) shown to be associated with BMI from a large-scale GWAS[18]. The independent sample of 418 individuals was used in the RbG study, based on the lower and upper ∼30% of a continuous genome-wide GRS distribution for BMI, constructed on the basis of results from a GWAS of BMI[19]. A total of 191 were within the lower tail and 227 were in the upper tail. The number of individuals with available data on the exposure (BMI) and cardiovascular outcomes are also presented*.

### Genotyping

Participants were genotyped using the Illumina HumanHap550 quad genome-wide SNP genotyping platform[20]. Participants were excluded due to having at least one of: incorrectly recorded sex, minimal or excessive heterozygosity, disproportionate levels of individual missingness, evidence of cryptic relatedness or non-European ancestry[20, 21]. SNPs with a minor allele frequency (MAF) of <1% and call rate of < 95% were removed and only SNPs that passed an exact test of Hardy-Weinberg equilibrium (*P*<5×10^−7^) were included. Imputation of genotypes was conducted with MACH 1.0.16 Markov Chain Haplotyping software, using CEPH individuals from phase 2 of the HapMap project as a reference (release #22).

### Measures of adiposity at age 17 and 21

At both ages, height was measured to the nearest centimetre using a stadiometer (SECA 213, Birmingham, UK) and weight to the nearest 0.1 kilogram, unshod and in light clothing, using electronic weighing scales (Marsden M-110, Rotherham, UK). BMI was calculated as weight (kg) divided by height-squared (m^2^).

### Cardiovascular phenotypes at age 17

The following cardiovascular phenotypes were used in MR analyses to assess the causal role of BMI on cardiovascular health at age 17.

#### Blood pressure and heart rate

Sitting BP and heart rate were measured in both arms with an Omron 705 IT oscillometric BP monitor in accordance with European Society of Hypertension guidelines[22]. The average of the final two of three readings was used and the arm with the greatest number of valid observations was used for analyses. From the systolic and diastolic BP readings (SBP and DBP, respectively), mean arterial pressure (MAP) was calculated (DBP + (SBP-DBP)/3). Pulse pressure (PP) was calculated as the difference between SBP and DBP.

#### Pulse-wave velocity (PWV)

Aortic stiffness (carotid-femoral PWV) was assessed using a Vicorder device (Skidmore Medical, UK). Participants rested supine on a couch with their head raised to 30°. Real-time pulse-wave measures were recorded between proximal (right carotid) and distal (the upper right thigh) sensor cuffs and with the time delay between the two simultaneously measured cardiac cycles measured. Transit distance was measured from suprasternal notch directly to the top of the thigh cuff. Measurements were taken until pressure waveforms over the carotid and thigh area were of high quality and reproducible. Three carotid-femoral PWV measurements, within ≤0.5 m/s of each other, were averaged.

#### Carotid intima-media thickness (cIMT)

Common carotid artery B-mode ultrasound images were acquired in the ear–to-ear plain with the head rotated to 45° from the midpoint using a Zonare Z.OneUltra system equipped with a L 10-5 linear transducer (Zonare Medical Systems, CA, US). Images were recorded in Digital Imaging and Communications in Medicine (DICOM) format as 10 second cine-loop files for offline analysis using the Carotid Analyzer (Medical Imaging Applications, Coralville, IA). Left and right cIMT were taken to be the average of three end-diastolic measurements located on the far-wall of a single segment of arterial wall 5−10 mm in length and 10 mm proximal to the bifurcation. The mean of left and right cIMT was calculated and used in analyses.

#### Left ventricular mass (LVM)

A sub-sample of study participants from the 17-year clinic underwent echocardiography using a HDI 5000 ultrasound machine (Phillips) and P4-2 Phased Array ultrasound transducer using a standard examination protocol. Left ventricular mass (LVM) was estimated according to American Society of Echocardiography (ASE) guidelines[23].

### Cardiovascular phenotypes at age 21

The following detailed cardiovascular phenotypes were also measured in the two RbG groups at age 21.

#### Blood pressure and heart rate

BP and heart rate were measured in the right arm in the supine position using a digital automated sphygmomanometer (Omron M6, Omron Healthcare, Netherlands) and the mean of two values used for analyses. MAP and PP were calculated in the same way as at 17 years.

#### Pulse-wave velocity

Arterial stiffness was assessed using applanation tonometry (SphygmoCor Vx, AtCor Medical, NSW, Australia), with PWV estimated using ECG-gated pulse waves travelling between carotid-femoral sites. The patient was rested in a supine position and a handheld tonometer was placed over the left carotid artery in order to allow the recording of 10-12 clear and reproducible pressure waveforms. The same tonometer was then used to measure a similar number of femoral arterial pulse waveforms in the inguinal crease at the top of the right leg. Transit distance was measured between the upper edge of the suprasternal notch and the femoral pulse measurement site via the umbilicus using a tape measure. The device software calculated the mean transit time (in milliseconds) from the recorded pulse waveforms and PWV was calculated as the transit distance/transit time.

#### Carotid intima-media thickness

Ultrasound assessment of the thickness of intima media interface was measured using an identical protocol to that described for age 17.

#### Cardiovascular Magnetic Resonance Imaging (MRI)

All measures were made using a 1.5T MR scanner (Avanto, Siemens Medical Solutions, Erlangen, Germany). Endocardial borders of the left ventricle (LV) were traced manually on short axis stacks at end-diastole and end-systole to evaluate end-diastolic volume (EDV) and end-systolic volume (ESV). Stroke volume (SV) was obtained by subtracting ESV from EDV. Epicardial borders were traced in end-diastole to calculate an epicardial volume. The EDV was subtracted from this volume, multiplied by assumed myocardial density to obtain LVM.

Flow quantification was performed through-plane in a cross-section of the ascending aorta as it passes the bifurcation of the pulmonary arteries using an ECG-gated spiral phase-contrast MR sequence, as described previously[24]. This technique allows images to be acquired within a short breath-hold (0.5 seconds) with a spatial resolution of 1.6 × 1.6 mm and a temporal resolution of 30 milliseconds. All images were processed using in-house plug-ins for the Open source software OsiriX (OsiriX Foundation, Geneva, Switzerland). Flow images were manually segmented (using the modulus images) and SV (ml) was measured and cardiac output (CO, L/min) was calculated as SV x heart rate. At the time of flow imaging, BP was simultaneously measured using MRI-compatible oscillometric sphygmomanometer (Datex Ohmeda). Systemic vascular resistance (SVR; measured in mmHg.L^−1^.min^−1^) was calculated by dividing the measured mean BP by CO. Total arterial compliance (TAC) was calculated by optimisation of the two-element Windkessel model, as previously described[25]. Briefly, the flow curves and SVR were used as inputs to the model. PP was calculated for a series of modelled pressure curves generated using a range of TAC values from 0.1 to 5.0 mL.mmHg^−1^ in increments of 0.01. The compliance value that gave the smallest error between the modelled PP and the true PP was taken to be the true compliance.

### Confounders

Due to potentially confounding effects, the following variables were added as covariates in the multivariable regression analyses: maternal education and household occupation, plus current smoking status and the most recent records of physical activity and, where available, dietary intake.

At enrolment, mothers reported their educational attainment and both her and her partner’s occupation. Highest household occupation was used to assign participants a household social class, using the 1991 British Office of Population Census Statistics classification[26]. Both maternal education and household social class were used as a general indication of socioeconomic position.

The most recent measurement of dietary intake of the participant was total energy intake at age 13, previously estimated from linear spline multi-level models of the combination of food frequency questionnaires and diet diaries[27]. Additionally, the most recent record of physical activity was defined as the counts per minute (CPM) and minutes spent in moderate-to-vigorous activity (MVPA) obtained from a subsample of participants at the 15-year clinic by an MTI Actigraph AM7164 2.2 accelerometer worn for 7 days[28]. Smoking status was obtained by questionnaire and participants were coded as “ever” or “never” smokers.

For analyses using data from the 21-year RbG group, physical activity and smoking status were obtained via questionnaire at the time of cardiovascular phenotyping; however, information on recent dietary intake was not available. Participants at this age were similarly classed as “ever” or “never” smokers and weekly exercise was categorised as either “never/rarely/<2 times a week” or “≥2 times a week”.

## STATISTICAL ANALYSES

### Pre-analysis transformations and adjustments

As the distribution of residuals from the linear regression of BMI on carotid-femoral PWV was positively skewed, values of these variables were log transformed. LVM measured at each age was indexed to height to the power of 2.7 (LVMI)[23]. To assess the impact of adiposity on central vascular measures over and above that caused by stature[29], SVR and TAC were adjusted for height using SVR × height^1.83^, TAC/height^1.83^[30], and both CO and SV measures were indexed to height by dividing CO by height^1.83^ and SV by height^2.04^. The use of “positive” and “inverse” throughout the text refer to directional association rather than clinical implication. Stata 14 (Stata Corp, Texas) and R (https://cran.r-project.org/) were used for all analyses.

### Multivariable regression

Observational associations between BMI and each cardiovascular phenotype at age 17 were assessed using multivariable linear regression in three models: (i) unadjusted, (ii) adjusted for age, sex, smoking status and dietary intake of the participant, household social class and maternal education and (iii) additionally adjusted for physical activity (added as a separate model due to sample size). Associations of the confounders with BMI, cardiovascular measures and weighted GRSs at each age were tested using linear regression.

### Mendelian randomization

The externally weighted GRS used as an instrument for BMI in MR analyses was generated from 97 independent SNPs shown to be reliably associated with BMI in the GWAS conducted by the Genetic Investigation of ANthropometric Traits (GIANT) consortium[18]. To generate the GRS, the dosage of each BMI-increasing allele at each locus in ALSPAC was weighted by the external effect size of the variant in the GWAS results[31, 32]. The doses were then added together and multiplied by the average external effect size of all the SNPs on BMI to reflect the number of average BMI-increasing alleles carried by each individual.

Two-stage least squares (2SLS) analysis was performed using the GRS as an instrument for BMI at age 17 (*ivreg2* command in Stata). *F*-statistics for the first-stage regression between the GRS and BMI were examined to check that the instruments were valid, satisfying the assumption that the instrument was sufficiently associated with the exposure[33]. The Durbin-Wu-Hausman (DWH) test for endogeneity was used to compare multivariable regression and IV effect estimates (*ivendog* command in Stata)[34].

### Recall-by-genotype

Linear regression was used to assess the association of the genome-wide GRS group allocation (upper vs. lower ∼30% of the genome-wide GRS distribution, as described above) with BMI and each of the cardiovascular phenotypes measured at age Each estimate therefore represents the mean difference in each variable with the corresponding mean difference in BMI between RbG groups.

### Sensitivity analyses

Both BP and heart rate are correlated with other cardiovascular measures, namely PWV, cIMT and LVMI[35-37]. To assess the causal association between BMI and these cardiovascular measures, independent of BP and heart rate, we took the residuals of the regression between each of these variables and both BP and heart rate and repeated MR and RbG main analyses using these residuals.

Additionally, evidence suggests that some of the cardiovascular phenotypes used in these analyses are not independent of height[23, 38, 39]. To account for this and the inconsistent residual correlation between BMI and height throughout the lifecourse[40], we assessed the association between the weighted GRS on height at age 17 and explored the impact of adjustment for height and height-squared on the association between the weighted GRS and BMI. We also adjusted both multivariable regression and MR analyses for height and height-squared measured at the same time of the BMI exposure and cardiovascular phenotype and compared these to the main analyses.

The use of multiple alleles in MR analyses increases the potential for unbalanced pleiotropic effects due to aggregation of invalid genetic instruments having an effect in one particular direction[13, 32]. To investigate the validity of the weighted GRS as an IV, the MR-Egger[41] approach was used to detect and accommodate violations of the MR assumptions, where the intercept of the MR-Egger test can be interpreted as an estimate of the average pleiotropic effect across the genetic variants, with a non-zero intercept term indicating overall directional pleiotropy. MR-Egger estimates were compared to those obtained from the inverse-variance weighted (IVW)[41, 42] and weighted median methods[43], which provide estimates of the causal effect of BMI on cardiovascular phenotypes under varying assumptions of instrument validity. As in the main analyses, the estimates of the association between each SNP and BMI were obtained from an independent external source, as to not induce weak instrument bias in a two-sample MR setting[44, 45].

Consistent with previous studies[46], we performed a sensitivity analysis using a weighted GRS that was limited to the genetic variants that were associated with BMI in the analysis of all people of European descent and excluded those that only reached genome-wide significance in only one sex or stratum (n=77) in the GIANT consortium[18]. Additionally, a previous study in a large sample based in the UK[46] suggested exclusion of three variants owing to pleiotropy (rs11030104, rs13107325 and rs3888190) and three SNPs that are not in Hardy-Weinberg equilibrium (P<1×10^−6^; rs17001654, rs2075650 and rs9925964). Therefore, as a sensitivity analysis, we excluded these additional SNPs, resulting in an instrument consisting of 71 independent SNPs.

Additionally, to assess the validity of the genome-wide GRS (derived from the results of the BMI GWAS conducted by Speliotes *et al*.[19] and used in main RbG analyses), MR analyses were conducted using the same genome-wide GRS as an instrument for BMI, scaled to represent the same difference in mean BMI per unit increase in the genome-wide GRS compared to the Locke *et al*. score, comprising 97 SNPs, used in main MR analyses.

### In silico recall-by-genotype analyses

To support the main RbG analyses at age 21 and the use of a genome-wide GRS, we conducted two *in silico* RbG analyses with available cardiovascular phenotypes at age 17. From the larger sample of individuals originally used in the MR analyses who had no missing data on BMI or cardiovascular outcomes at age 17 (N=1190), we randomly sampled 200 individuals from the lower and upper ∼30% of the distribution of i) the Locke *et al*. GRS (N=97 SNPs, constructed using external weighting)[18] and ii) the genome-wide GRS from the Speliotes *et al*. GWAS used in main RbG analyses[19]. To do this, each of the scores was sorted by value and 400 individuals were kept from both the lower and upper ∼30% of the distribution. Of those, 50% were randomly selected for analyses, leaving 200 at each of the lower and upper tails (comparable to the selection criteria for the main RbG study). This was also tested with an iterative random sampling approach (results not shown).

## RESULTS

The MR cohort were 17.8 years old (SD = 0.4), consisted of 47.8% females and had an average BMI of 22.7kg/m^2^ (SD = 4) (Table 1). In the RbG study, individuals were 21.5 years old (SD = 0.9), had an average BMI of 24.5kg/m^2^ (SD = 5.7) and 65.8% were females (Table 2).

**Table 1.** Descriptive statistics for ALSPAC 17-year clinic.

| Variable | N | Mean (SD) or percentage |
| --- | --- | --- |
| *Participant's phenotypes* |  |  |
| Age (years) | 3493 | 17.79 (0.42) |
| Sex (% female) | 7909 | 47.81 |
| BMI (kg/m<sup>2</sup>) | 3404 | 22.73 (3.99) |
| SBP (mmHg) | 3172 | 118.61 (10.95) |
| DBP (mmHg) | 3172 | 63.65 (6.56) |
| PP (mmHg) | 3172 | 54.96 (10.08) |
| MAP (mmHg) | 3172 | 81.97 (6.78) |
| Mean cIMT (mm) | 3143 | 0.48 (0.04) |
| Carotid-femoral PWV (m/s) | 2529 | 5.75 (0.69) |
| LVMI (g/m<sup>2.7</sup>) | 1420 | 28.93 (6.11) |
| Heart rate (bpm) | 3172 | 64.45 (9.80) |
| Smoking status (% ever smoked) | 2844 | 51.55 |
| *Physical activity* |  |  |
| CPM (counts) | 1687 | 484.09 (180.99) |
| MVPA (minutes) | 1687 | 23.73 (18.81) |
| Dietary intake (kcal) | 7141 | 2260.24 (184.61) |
| *Parental phenotypes* |  |  |
| Highest household social class | 6598 |  |
| I | 947 | 14.35 |
| II | 2902 | 43.98 |
| III (non-manual) | 1651 | 25.02 |
| III (manual) | 769 | 11.66 |
| IV | 290 | 4.40 |
| V | 39 | 0.59 |
| Maternal education | 6982 |  |
| CSE | 1156 | 16.56 |
| Vocational | 635 | 9.09 |
| O-Level | 2453 | 35.13 |
| A-Level | 1700 | 24.35 |
| Degree | 1038 | 14.87 |

ALSPAC = Avon Longitudinal Study of Parents and Children; BMI = body mass index; BP = blood pressure; cIMT = carotid intima-media thickness; CSE = certificate of secondary education; DBP = diastolic blood pressure; LVMI = left ventricular mass indexed to height<sup>2.7</sup>; MAP = mean arterial pressure; PWV = pulse wave velocity; SBP = systolic blood pressure; SD = standard deviation

**Table 2.** Descriptive statistics for ALSPAC 21-year RbG group.

| Variable | N | Mean (SD) or percentage |
| --- | --- | --- |
| *Participant's phenotype* |  |  |
| Age (years) | 418 | 21.51 (0.94) |
| Sex (% female) | 418 | 65.79 |
| BMI (kg/m<sup>2</sup>) | 418 | 24.52 (5.70) |
| SBP (mmHg) | 415 | 116.97 (10.28) |
| DBP (mmHg) | 415 | 67.23 (6.64) |
| PP (mmHg) | 415 | 49.74 (9.57) |
| MAP (mmHg) | 415 | 83.81 (6.65) |
| Mean cIMT (mm) | 417 | 0.46 (0.04) |
| Carotid-femoral PWV (m/s) | 404 | 5.58 (0.74) |
| SVR (mmHg/L/min) | 386 | 15.62 (3.02) |
| TAC (mL/mmHg) | 386 | 1.02 (0.23) |
| LVMI (g/m<sup>2.7</sup>) | 386 | 21.42 (4.15) |
| Heart rate (bpm) | 406 | 61.24 (9.43) |
| SV (ml) | 386 | 89.69 (16.96) |
| CO (l/min) | 386 | 5.70 (1.16) |
| Smoking status (% ever smoked) | 417 | 17.27 |
| Weekly exercise (% never/rarely/<2 per week) | 299 | 38.46 |
| *Parental phenotypes* |  |  |
| Highest household social class | 370 |  |
| I | 91 | 24.59 |
| II | 169 | 45.68 |
| III (non-manual) | 76 | 20.54 |
| III (manual) | 27 | 7.30 |
| IV | 7 | 1.89 |
| Maternal education | 380 |  |
| CSE | 31 | 8.16 |
| Vocational | 28 | 7.37 |
| O-Level | 109 | 28.68 |
| A-Level | 120 | 31.58 |
| Degree | 92 | 24.21 |

ALSPAC = Avon Longitudinal Study of Parents and Children; BMI = body mass index; BP = blood pressure; cIMT = carotid intima-media thickness; CO = cardiac output; CSE = certificate of secondary education; DBP = diastolic blood pressure; LVMI = left ventricular mass indexed to height<sup>2.7</sup>; MAP = mean arterial pressure; PP = pulse pressure; PWV = pulse wave velocity; SBP = systolic blood pressure; SD = standard deviation; SV = stroke volume; SVR = systemic vascular resistance; TAC = total arterial compliance

### Confounder analyses

BMI and all of the cardiovascular phenotypes were associated with a majority of the confounding factors (Supplementary Tables S1 and S2, respectively).

### Multivariable regression

After adjusting for potential confounders, multivariable regression analyses provided evidence for positive associations of measured BMI with SBP, DBP, PP, MAP, LVMI and heart rate at age 17 (Table 3), as well as an inverse association with carotid-femoral PWV.

**Table 3.** Multivariable regression associations between BMI and 436 cardiovascular phenotypes in ALSPAC 17-year clinic.

| Outcome (units) | N | Difference in mean outcome per 1kg/m<sup>2</sup> higher BMI (95% CI) | P-value | N | Difference in mean outcome per 1kg/m<sup>2</sup> higher BMI (95% CI)<sup>1</sup> | P-value | N | Difference in mean outcome per 1kg/m<sup>2</sup> higher BMI (95% CI)<sup>2</sup> | P-value |
| --- | --- | --- | --- | --- | --- | --- | --- | --- | --- |
| SBP (mmHg) | 3108 | 0.86 (0.76, 0.95) | 7.43x10<sup>-74</sup> | 2389 | 0.84 (0.74, 0.94) | 6.31x10<sup>-64</sup> | 1033 | 0.83 (0.68, 0.98) | 1.00x10<sup>-25</sup> |
| DBP (mmHg) | 3108 | 0.52 (0.47, 0.58) | 4.67x10<sup>-77</sup> | 2389 | 0.49 (0.42, 0.56) | 2.73x10<sup>-44</sup> | 1033 | 0.47 (0.36, 0.58) | 1.44x10<sup>-17</sup> |
| PP (mmHg) | 3108 | 0.33 (0.25, 0.42) | 1.98x10<sup>-13</sup> | 2389 | 0.35 (0.26, 0.44) | 6.08x10<sup>-15</sup> | 1033 | 0.36 (0.22, 0.49) | 3.97x10<sup>-07</sup> |
| MAP (mmHg) | 3108 | 0.63 (0.58, 0.69) | 1.13x10<sup>-109</sup> | 2389 | 0.61 (0.54, 0.68) | 2.89x10<sup>-68</sup> | 1033 | 0.59 (0.48, 0.69) | 5.64x10<sup>-27</sup> |
| Mean cIMT (mm) | 3079 | 0.0001<br>(0.0003, 0.0005) | 0.72 | 2368 | -0.0001<br>(-0.001, 0.0004) | 0.65 | 1028 | 0.0003<br>(-0.0005, 0.001) | 0.48 |
| Carotid-femoral PWV (log(m/s)) | 2495 | -0.001<br>(-0.002, 0.0004) | 0.20 | 1957 | -0.001<br>(-0.002, 0.001) | 0.20 | 867 | -0.003<br>(-0.005, -0.0004) | 0.02 |
| LVMI (g/m<sup>2.7</sup>) | 1420 | 0.79 (0.72, 0.86) | 7.66x10<sup>-108</sup> | 1151 | 0.83 (0.75, 0.91) | 2.02x10<sup>-99</sup> | 569 | 0.95 (0.83, 1.07) | 5.40x10<sup>-57</sup> |
| Heart rate (bpm) | 3108 | 0.21 (0.12, 0.29) | 3.50x10<sup>-06</sup> | 2389 | 0.20 (0.10, 0.31) | 0.0001 | 1033 | 0.24 (0.08, 0.39) | 0.003 |

ALSPAC = Avon Longitudinal Study of Parents and Children; BMI = body mass index; BP = blood pressure; CI = confidence interval; cIMT = carotid intima-media thickness; DBP = diastolic blood pressure; LVMI = left ventricular mass indexed to height<sup>2.7</sup>; MAP = mean arterial pressure; PP = pulse pressure; PWV = pulse wave velocity; SBP = systolic blood pressure
<sup>1</sup>Adjusted for age, gender, smoking and dietary intake of the participant and maternal education and household social class
<sup>2</sup>Additionally adjusted for physical activity of the participant

### Mendelian randomization

Each allele increase in the weighted GRS (comprising 97 SNPs) was associated with a 0.12kg/m^2^ (95% CI: 0.10, 0.14; *P*=9.53×10^−28^) higher BMI, explaining 3% of the variance (Figure 3). Unlike the direct measure of BMI and the cardiovascular phenotypes, the GRS was not associated with a majority of confounders (Supplementary Table S3).There was evidence for a positive effect of each kg/m^2^ higher BMI on SBP (difference: 0.79mmHg; 95% CI: 0.30, 1.28; P=0.002), DBP (difference: 0.29mmHg; 95% CI: 0.0002, 0.59; *P*=0.05), PP (difference: 0.49mmHg; 95% CI: 0.03, 0.96; *P*=0.04), MAP (difference: 0.46mmHg; 95% CI: 0.16, 0.75; *P*=0.002) and LVMI (difference: 1.07g/m^2.7^; 95% CI: 0.62, 1.52;P=3.87×10^−06^) (Table 4). F-statistics for these analyses ranged from 36 to 123. There was no strong evidence that the results from MR analyses were different to those from the multivariable regression analyses (all *P*-values for comparison > 0.12).

**Figure 3.**
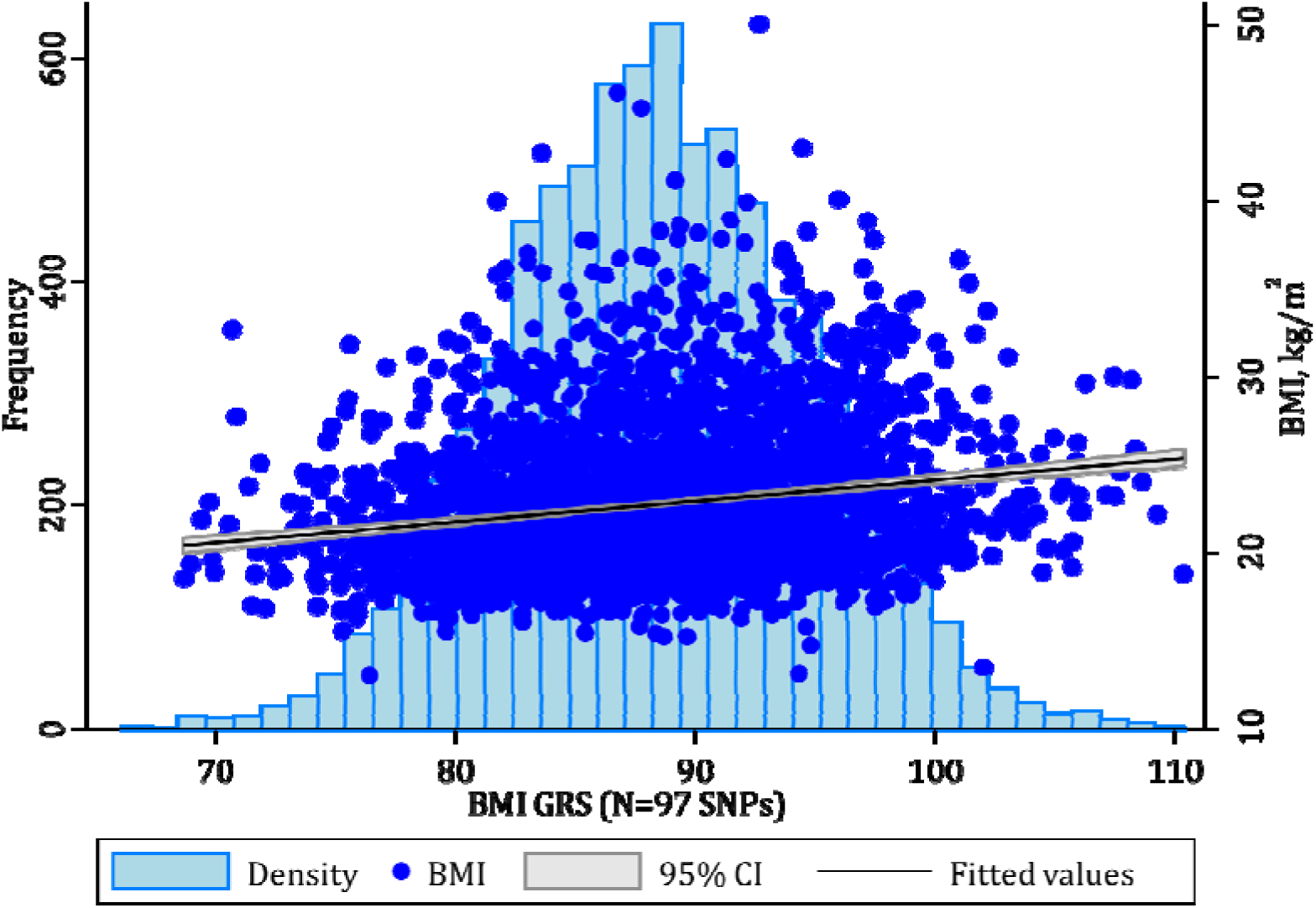
Association between weighted GRS (comprising 97 SNPs) and BMI in ALSPAC 17-year clinic. *The light blue histogram represents the weighted GRS (comprising 97 SNPs) distribution (frequency, left-hand axis). The mid-blue scatter plot and linear trend with corresponding 95% confidence intervals represent the association between the same weighted GRS and BMI (kg/m^2^, right-hand axis)*.

**Table 4.** Mendelian randomization analyses of the association between BMI and cardiovascular phenotypes in ALSPAC 17-year clinic.

| Outcome (units) | N | Difference in mean outcome per 1kg/m<sup>2</sup> higher BMI (95% CI) | P-value | F-stat | P-value for difference in multivariable regression and MR analyses<sup>1</sup> |
| --- | --- | --- | --- | --- | --- |
| SBP (mmHg) | 3108 | 0.79 (0.30, 1.28) | 0.002 | 115.76 | 0.78 |
| DBP (mmHg) | 3108 | 0.29 (0.0002, 0.59) | 0.05 | 115.76 | 0.12 |
| PP (mmHg) | 3108 | 0.49 (0.03, 0.96) | 0.04 | 115.76 | 0.49 |
| MAP (mmHg) | 3108 | 0.46 (0.16, 0.75) | 0.002 | 115.76 | 0.24 |
| Mean cIMT (mm) | 3079 | 0.002 (-0.001, 0.004) | 0.14 | 122.97 | 0.15 |
| Carotid-femoral PWV (log(m/s)) | 2495 | -0.001 (-0.01, 0.01) | 0.74 | 86.30 | 0.92 |
| LVMI (g/m<sup>2.7</sup>) | 1420 | 1.07 (0.62, 1.52) | 3.87x10<sup>-06</sup> | 36.24 | 0.21 |
| Heart rate (bpm) | 3108 | -0.05 (-0.51, 0.41) | 0.82 | 115.76 | 0.26 |

*ALSPAC = Avon Longitudinal Study of Parents and Children; BMI = body mass index; BP = blood pressure; CI = confidence interval; cIMT =* *carotid intima-media thickness; DBP = diastolic blood pressure; LVMI = left ventricular mass indexed to height<sup>2.7</sup>; MR = Mendelian* *randomization; MAP = mean arterial pressure; PP = pulse pressure; PWV = pulse wave velocity; SBP = systolic blood pressure; SNP = single* *nucleotide polymorphism*
<sup>1</sup>*P-value obtained from Durbin-Wu-Hausman test of heterogeneity*

### Recall-by-genotype

Difference in mean BMI between RbG groups was 3.85kg/m^2^ (95% CI:2.53, 4.63; *P*=6.09×10^−11^) (Table 5, Figure 4). Measures of both BMI and cardiovascular outcomes were associated with a majority of confounders when assessed as a whole sample (Supplementary Tables S4 and S5, respectively). There was no strong evidence that the RbG group allocation was associated with confounders (Supplementary Table S6).

**Figure 4.**
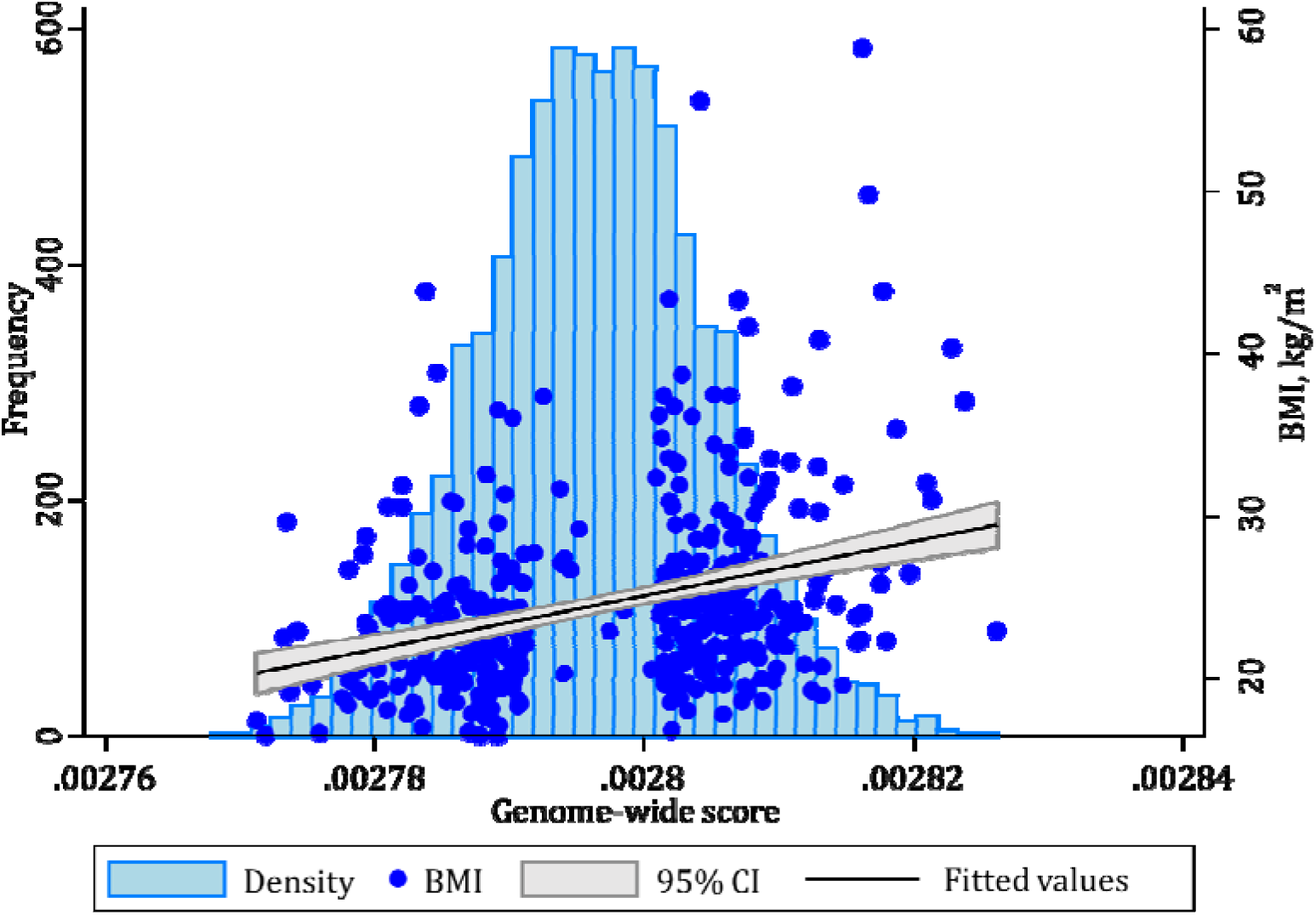
Association between RbG 486 groups and BMI in ALSPAC 21-year RbG group. *The light blue histogram represents the genome-wide GRS distribution (frequency, left-hand axis). The mid-blue scatter plot and linear trend with corresponding 95% confidence intervals represent the association between the same genome-wide GRS and BMI (kg/m^2^, right-hand axis) included in the RbG groups*.

**Table 5.** Association between RbG groups and cardiovascular measures in ALSPAC 21-year RbG group

| Outcome (units) | N | Difference in mean outcome per 3.58kg/m<sup>2</sup> higher BMI (95% CI) | P-value |
| --- | --- | --- | --- |
| BMI (kg/m<sup>2</sup>) | 418 | 3.58 (2.53, 4.63) | $6.09 \times 10^{-11}$ |
| *Cardiovascular outcomes* |  |  |  |
| SBP (mmHg) | 415 | 3.70 (1.74, 5.66) | 0.0002 |
| DBP (mmHg) | 415 | 2.25 (0.98, 3.52) | 0.001 |
| PP (mmHg) | 415 | 1.45 (-0.40, 3.30) | 0.12 |
| MAP (mmHg) | 415 | 2.73 (1.47, 3.99) | 0.00003 |
| Mean cIMT (mm) | 417 | 0.001 (-0.01, 0.01) | 0.82 |
| Carotid-femoral PWV (log(m/s)) | 404 | 0.03 (0.01, 0.06) | 0.01 |
| SVR (mmHg/L/min.m<sup>1.83</sup>) | 386 | -1.03 (-2.49, 0.43) | 0.17 |
| TAC (mL/mmHg/m<sup>1.83</sup>) | 386 | 0.01 (-0.01, 0.02) | 0.37 |
| LVMl (g/m<sup>2.7</sup>) | 386 | 1.65 (0.83, 2.47) | 0.0001 |
| Heart rate (bpm) | 406 | -0.08 (-1.93, 1.77) | 0.93 |
| SV (ml/m<sup>2.04</sup>) | 386 | 1.49 (0.62, 2.35) | 0.001 |
| CO (l/min/m<sup>1.83</sup>) | 386 | 0.11 (0.03, 0.19) | 0.01 |

ALSPAC = Avon Longitudinal Study of Parents and Children; BMI = body mass index; BP = blood pressure; CI = confidence interval; cIMT = carotid intima-media thickness; CO = cardiac output; DBP = diastolic blood pressure; LVMl = left ventricular mass indexed to height<sup>2.7</sup>; MAP = mean arterial pressure; PP = pulse pressure; PWV = pulse wave velocity; SBP = systolic blood pressure; SV = stroke volume; SVR = systemic vascular resistance; TAC = total arterial compliance

Of the cardiovascular measures that overlapped between the two methods (MR and RbG), the RbG groups were associated with higher SBP (difference in mean between higher vs. lower RbG groups: 3.70mmHg; 95% CI:1.74, 5.66; *P*=0.0002), DBP (difference: 2.25mmHg; 95% CI: 0.98, 3.52; *P*=0.001), MAP (difference: 2.73mmHg; 95% CI: 1.47, 3.99; *P*=0.00003) and carotid-femoral PWV (difference: 0.03log(m/s); 95% CI: 0.01, 0.06; *P*=0.01) (Table 5). Scaling the effect estimates to represent a kg/m^2^ higher BMI, as in the MR analyses, these results are equivalent to a 1.03mmHg higher SBP, a 0.63mmHg higher DBP, a 0.76mmHg higher MAP and a 0.01log(m/s) higher carotid-femoral PWV. There was therefore considerable consistency between effect estimates on the overlapping phenotypes at both ages, i.e. each kg/m^2^ higher BMI had a causal effect of similar magnitude on SBP, DBP and MAP, whilst showing no association with heart rate or cIMT (Figure 5). However, there was evidence for a positive causal effect of BMI on carotid-femoral PWV in RbG analyses at age 21 that was not evident in MR analyses at 17 years.

**Figure 5.**
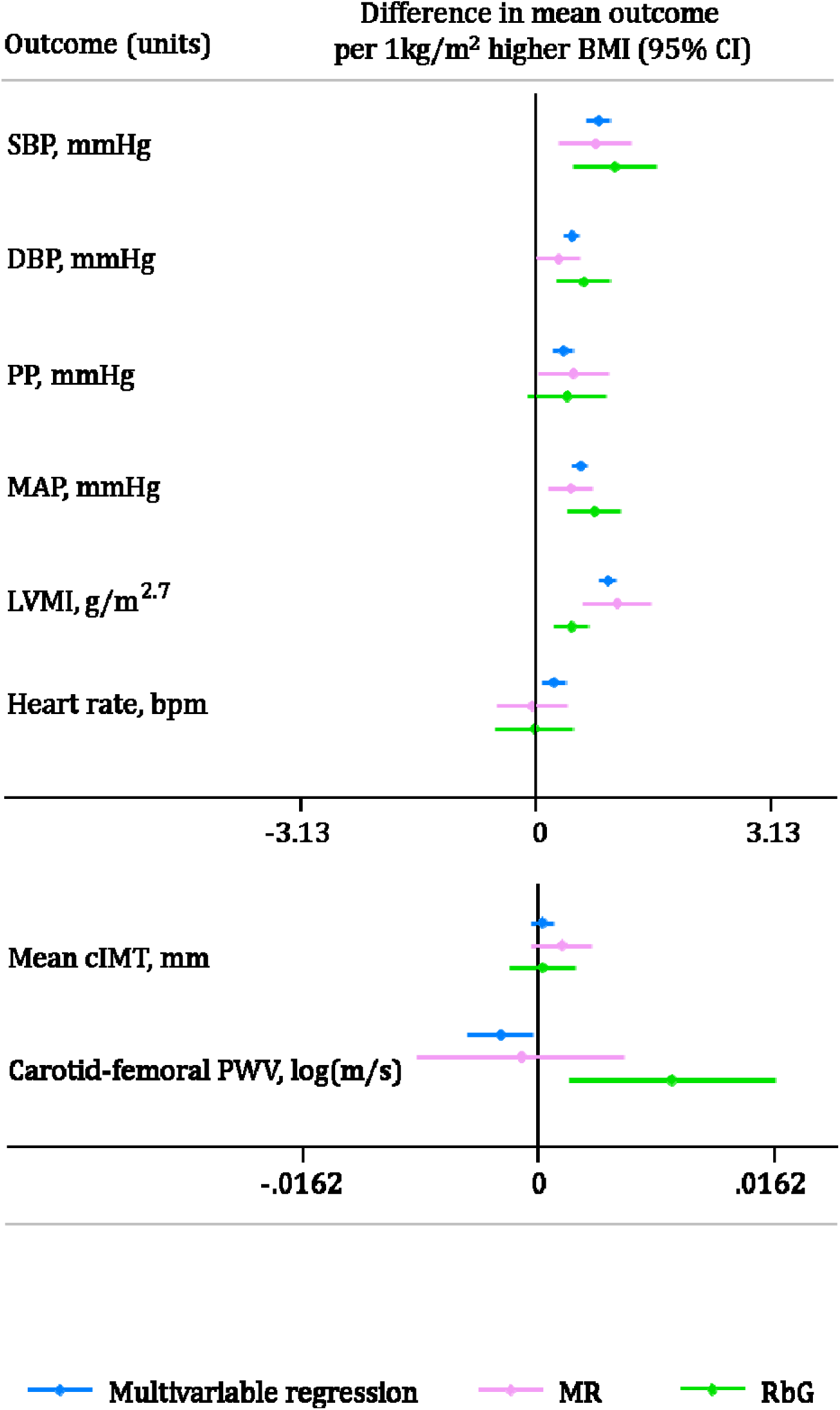
Comparison of estimates from multivariable regression, MR and RbG methodologies for the difference of all cardiovascular phenotypes available at both ages (graphs are separated by scale similarities) per 1kg/m^2^ higher BMI.

In addition to these cardiovascular measures, the RbG framework enabled the collection of more precise cardiovascular phenotypes and showed a positive causal role of higher BMI on MRI-derived LVMI (difference in mean between higher vs. lower RbG groups: 1.65g/m^2.7^; 95% CI: 0.83, 2.47; *P*=0.0001), SV (difference: 1.49ml/m^2.04^; 95% CI: 0.62, 2.35; *P*=0.001) and CO (difference: 0.11l/min/m^1.83^; 95%CI: 0.03, 0.20; *P*=0.01), with no evidence of a difference in SVR or TAC.

### Sensitivity analyses

After adjusting for SBP and heart rate, both MR and RbG results for the effect of BMI on cIMT, carotid-femoral PWV and LVMI were mostly consistent with main analysis (Supplementary Table S7). The one exception was the positive effect of BMI on carotid-femoral PWV shown in RbG analysis, which attenuated to the null following adjustment for SBP and heart rate (estimate: 0.02(log(m/s)); 95% CI:-0.01, 0.04; *P*=0.18).

The weighted GRS (comprising 97 SNPs), used in MR analyses, was not associated with height or height-squared at age 17. Adjusting for both height and height-squared made no difference to the association between the GRS and BMI at age 17 (Supplementary Table S8a), multivariable regression analyses (Supplementary Table S8b) or MR analyss (Supplementary Table S8c).

Where pleiotropy is perfectly balanced, an informative GRS is sufficient in an MR analysis, but this method is less able to cope with unbalanced pleiotropic effects (where invalid genetic instruments have an aggregate effect in one particular direction)[41]. The MR-Egger can go some way to accommodate violation of MR assumptions through pleiotropy, where the intercept term provides an estimate for unbalanced pleiotropy. This test provided no evidence for unbalanced pleiotropic effects of the genetic variants included within the GRS on any cardiovascular outcome (all *P*-values for the intercept ≥ 0.24) (Supplementary Table S9). The MR effect estimates from IVW, MR-Egger and weighted median analyses for the causal effect of BMI on the cardiovascular phenotypes were largely consistent with the main analyses, though the effect estimate of BMI on SBP did not agree for the weighted median analyses compared to the main MR, IVW and MR-Egger estimates, albeit with very wide confidence intervals (Supplementary Table S9 and Supplementary Figures S1a-S1e).

Both the instrument containing 77 SNPs (found in Europeans and analyses of both sexes only in the GIANT GWAS, Supplementary Figure S2) and 71 SNPs (found in Europeans only and removing potentially pleiotropic SNPs and those that were not in HWE in a large sample based in the UK, Supplementary Figure S3) were associated with BMI to a comparable extent as the GRS comprising the full set of 97 SNPs (Supplementary Table S10) and produced similar results to the main analyses (Supplementary Tables S11b and S11b, respectively). Similarly, when the genome-wide GRS initially used to recall individuals to the RbG study was implemented in MR analyses, the GRS was associated with a comparable change in BMI (Supplementary Table S10, Supplementary Figure S4) and produced similar results to main MR analyses (Supplementary Table S12).

### In silico recall-by-genotype analyses

In both *in silico* RbG analyses, the difference in mean BMI between the two RbG groups was similar (1.63kg/m^2^; 95% CI: 0.93, 2.32; *P*=5.89×10^−06^ and 1.65kg/m^2^; 95% CI: 0.91, 2.38; *P*=1.43×10^−05^ between RbG groups sampled from lower and upper ∼30% of the tails of the Locke *et al*. GRS and the Speliotes *et al*. genome-wide GRS distributions, respectively), each explaining 5% of the variance in BMI (Supplementary Table S13 and Supplementary Figures S5 and S6, respectively). Most results from both *in silico* RbG analyses were in the same direction of effect and also in the same direction as results from the main RbG analyses (Supplementary Table S14a and S14b, respectively). For example, scaling the effect estimates to represent a kg/m^2^ higher BMI, as in the main analyses, the equivalent effect estimates using the weighted GRS (comprising 97 SNPs) and the genome-wide GRS, respectively, were 0.87mmHg (95% CI:-1.23, 2.97; *P*=0.42) and 0.67mmHg (95% CI:-1.46, 2.79; *P*=0.54) for SBP, 1.45g/m^2.7^ (95% CI: 0.27, 2.63; *P*=0.02) and 1.83g/m^2.7^ (95% CI: 0.68, 2.98; *P*=0.002) for LVMI and -0.01log(m/s) (95% CI: -0.03, 0.02; *P*=0.59) and -0.01log(m/s) (95% CI: -0.03, 0.01; *P*=0.22) for carotid-femoral PWV. However, the direction of effect for DBP, MAP and heart rate differed between the two *in silico* sensitivity analyses (with one always being consistent with the main analyses), but these *in silico* analyses had wide confidence intervals. The iterative random sampling produced the same results (not shown).

## Discussion

### Summary and comparison of findings

In a large cohort of young adults, we employed two complementary analyses (MR and RbG) to investigate the causal effect of higher BMI on measures of cardiovascular structure and function and compared these to adjusted multivariable regression results. Alongside multivariable regression and MR analyses, the RbG method predominantly allowed the collection of extremely precise cardiovascular phenotypes that would otherwise be impractical at scale.

Regarding the cardiovascular phenotypes that were used across MR, RbG and multivariable regression analyses in the current study, our results are consistent with previous observational studies in children and adults[47-49] and across all three methodologies used. Results suggest that higher BMI causes higher BP (SBP, DBP, PP and MAP) and higher LVMI, the latter suggesting adverse cardiac structure, even in young adults. Furthermore, the similarity of findings across these methods, given different sources of bias between the MR and RbG on the one hand[41, 50] and multivariable regression on the other[13], strongly support causality in this instance. If further sustained through adulthood, these cardiovascular effects of higher BMI are likely to increase CVD risk and CVD-specific mortality in later life[51-55].

Previous multivariable regression results from smaller observational studies in children and adolescents have found higher BMI to be associated with faster PWV and thicker cIMT[56-59]. In contrast to this, our three methods gave results that did not support a causal effect for cIMT. This suggests that previous studies may have been influenced by residual confounding or bias, for which we have been better able to control here. This conclusion is also supported to some extent by analysis of the same variables at a lower age in an observational context[47]. With less consistent results across our analysis approaches, higher BMI was associated with slower PWV at age 17 years (i.e. healthier PWV) in multivariable regression analysis and a faster PWV (i.e. worse PWV) in RbG analysis, with MR analysis showing a null association (Figure 5). Indeed, another previous study suggests that the relationship between BMI and PWV may be inverse in youth and becomes positive at older ages[60].

As might be expected, higher BMI resulted in increased CO in our RbG study and, although contrary to other observational studies at this age[61, 62], this appeared to be solely driven by SV, as neither our MR or RbG analyses suggested a causal effect of BMI on heart rate. It is possible that previously reported associations between BMI and heart rate may be a result of unmeasured confounding. The BMI-mediated change in SV (and consequently CO) seen at age 21 is therefore likely to at least partially account for the cardiac hypertrophy and higher BP that we see in multivariable regression, MR and RbG both ages in the data analysed here.

### Strengths and limitations

A key strength is the comparison of results from confounder-adjusted multivariable regression, MR and RbG, along with a range of appropriate sensitivity analyses, which together provide strong evidence for a causal effect of higher BMI on BP and cardiac structure (LVMI) in young adulthood. This is one of the first uses of RbG for BP and cardiac structure, where we could compare results directly to multivariable regression and MR within the same general population. The consistency between results found from RbG with 418 participants to those from MR with 7,909 participants suggests this approach is valid and statistically efficient. Furthermore, the RbG method allowed the collection of precise cardiovascular phenotypes that would otherwise not have been possible in sample sizes required for MR. For example, we could explore the impact of BMI on SV and CO, measures that are prohibitively expensive to undertake in several thousands of participants.

In contrast to these strengths, it is of course the case that both the MR and RbG analyses may be biased if the IV analyses assumptions are violated[13]. These require, firstly, that the genetic instruments need to be robustly related to the exposure (here BMI). We used variants in both MR and RbG that have been shown to be genome-wide significant and replicated; the first-stage F-statistics, a measure of instrument strength, were high for all the MR analyses. Secondly, it is assumed that confounders of the observational BMI-cardiovascular outcome association are not related to the genetic instrument. There is empirical evidence that this is unlikely to be the case[63] and for observed confounders we demonstrate it in our analyses here. Thirdly, it is assumed that there is no path from the genetic instrument to the outcomes other than through BMI, which may result from horizontal pleiotropy. Using an aggregate allelic score increases the possibility of horizontal pleiotropy and we therefore undertook sensitivity analyses to explore this[41]. Across a range of sensitivity analysis (including the MR-Egger and weighted median approaches in the MR analyses, limiting the GRS to different subsets of genetic instruments and performing *in silico* RbG analyses using different genetic proxies), provided broadly similar results to main analyses, lending more confidence to the causal estimates and direction of effect with higher BMI and suggesting that these were not largely driven by horizontal pleiotropy.

Although the RbG approach enabled sampling from lower and upper ∼30% of a genome-wide GRS, which produced a difference of ∼3.5kg/m^2^ in BMI, the genome-wide nature of the score could be considered less refined than the GRS used in MR analyses, comprising 97 SNPs shown to be robustly associated with BMI in a large meta-analyses of GWASs[18]. Despite this, sensitivity analyses performed showed that the genome-wide GRS provided comparable results to MR-analyses and *in silico* RbG analyses.

One complication in some of the sensitivity analyses performed (specifically, adjusting for variables including BP, heart rate and height in multivariable and MR analyses) is the potential for inducing collider bias[64, 65]. However, due to the overall consistency in effect estimates generated from the various sensitivity analyses, this is unlikely to be the case in this study. More generally, it is possible that some of the differences in effect size of BMI on cardiovascular outcomes (for example, carotid-femoral PWV) between methodologies may relate to the difference in age at which the methods were applied. In addition, given the small range of some of the cardiovascular outcomes (for example, cIMT) in these young individuals and the potentially small effect size of BMI, power to detect such small effect sizes in this context may be limited.

Further, we adjusted for a range of potentially confounding factors in multivariable regression analyses but even in such a comprehensive longitudinal cohort, it can be difficult to accurately measure (and therefore appropriately account for) all confounders. Indeed, this illustrates the need for better methods (such as MR and RbG employed here) to assess the causal nature of the association between BMI and cardiovascular health.

## Conclusion

With this innovative study design, using complementary multivariable regression, MR and RbG analyses, together with a range of appropriate sensitivity analyses, we found results that strongly suggest a causal role of higher BMI resulting in adverse levels of BP and LVMI. These findings support efforts to reduce BMI and prevent the obesity burden from a young age, with the aim of attenuating the development of vascular and cardiac changes, known to be the precursors of long-term adverse cardiovascular outcomes, and preventing the development of additional peripheral vascular damage not yet evident at this early stage of life.

## Acknowledgements

We are extremely grateful to all the families who took part in this study, the midwives for their help in recruiting them, and the whole ALSPAC team, which includes interviewers, computer and laboratory technicians, clerical workers, research scientists, volunteers, managers, receptionists and nurses. We’d also like to thank Frank Dudbridge for ongoing conversations regarding genetic risk scores used in Mendelian randomization methodologies. The UK Medical Research Council (MRC) and the Wellcome Trust (Grant ref: 102215/2/13/2) and the University of Bristol provide core support for ALSPAC. This publication is the work of the authors and KHW and NJT serve as guarantors for the contents of this paper. This research was specifically funded through grants from the British Heart Foundation (grant numbers: RG/10/004/28240, PG/06/145 and CS/15/6/31468) and the UK MRC, University of Bristol and Wellcome Trust (grant numbers: MC_UU_12013/1-9, 096989/Z11/Z, 086676/7/08/Z, MR-M009351/1, MR/M020894/1 and 202802/Z/16/Z). GWAS data was generated by Sample Logistics and Genotyping facilities at Wellcome Sanger Institute and LabCorp (Laboratory Corporation of America) using support from 23andMe.

## References

1. Calle EE, Thun MJ, Petrelli JM, Rodriguez C, Heath CWJ. Body-Mass Index and Mortality in a Prospective Cohort of U.S. Adults. New England Journal of Medicine. 1999; 341(15): 1097–105.

2. Prospective Studies Collaboration. Body-mass index and cause-specific mortality in 900,000 adults: collaborative analyses of 57 prospective studies. The Lancet. 373(9669): 1083–96.

3. Nordestgaard BG, Palmer TM, Benn M, Zacho J, Tybjæsrg-Hansen A, Davey Smith G, et al. The Effect of Elevated Body Mass Index on Ischemic Heart Disease Risk: Causal Estimates from a Mendelian Randomisation Approach. PLOS Medicine. 2012; 9(5): e1001212.

4. Adams TD, Gress RE, Smith SC, Halverson RC, Simper SC, Rosamond WD, et al. Long-Term Mortality after Gastric Bypass Surgery. New England Journal of Medicine. 2007;357(8): 753–61.

5. Christou NV SJ, Liberman M, Look D, Auger S, McLean APH, MacLean LD. Surgery decreases long-term mortality, morbidity, and health care use in morbidly obese patients. Ann Surg. 2004; 240: 416–24.

6. Hägg S, Fall T, Ploner A, Mägi R, Fischer K, Draisma HH, et al. Adiposity as a cause of cardiovascular disease: a Mendelian randomization study. International Journal of Epidemiology. 2015; 44: 578–586.

7. Mozaffarian D, Benjamin EJ, Go AS, Arnett DK, Blaha MJ, Cushman M, et al. Heart Disease and Stroke Statistics—2015 Update. Circulation. 2015:131: e29–e322.

8. Halcox JPJ DJ. Childhood origins of endothelial dysfunction. Heart. 2005;91:1272–4.

9. Oren A, Vos LE, Uiterwaal CM, Grobbee DE, Bots ML. Cardiovascular risk factors and increased carotid intima-media thickness in healthy young adults: The atherosclerosis risk in young adults (arya) study. Archives of Internal Medicine. 2003; 163(15): 1787–92.

10. Wildman RP, Mackey RH, Bostom A, Thompson T, Sutton-Tyrrell K. Measures of Obesity Are Associated With Vascular Stiffness in Young and Older Adults. Hypertension. 2003; 42(4): 468.

11. Lorber R, Gidding SS, Daviglus ML, Colangelo LA, Liu K, Gardin JM. Influence of systolic blood pressure and body mass index on left ventricular structure in healthy African-American and white young adults: the CARDIA study. Journal of the American College of Cardiology. 2003; 41(6): 955–60.

12. Haycock PC, Burgess S, Wade KH, Bowden J, Relton C, Davey Smith G. Best (but oft-forgotten) practices: the design, analysis, and interpretation of Mendelian randomization studies. The American Journal of Clinical Nutrition. 2016:103: 965–978.

13. Davey Smith G HG. Mendelian randomization: genetic anchors for causal inference in epidemiology studies. Hum Mol Genet. 2014:23:R89–R98.

14. Ware JJ, Timpson N, Davey Smith G, Munafò MR. A recall-by-genotype study of CHRNA5-A3-B4 genotype, cotinine and smoking topography: study protocol. BMC Medical Genetics. 2014;15:13

15. Hellmich C, Durant C, Jones MW, Timpson NJ, Bartsch U, Corbin LJ. Genetics, sleep and memory: a recall-by-genotype study of ZNF804A variants and sleep neurophysiology. BMC Medical Genetics. 2015; 16: 96.

16. Boyd A GJ, Macleod J, Lawlor DA, Fraser A, Henderson J, Molloy L, Ness A, Ring S, Davey Smith G. Cohort Profile: the ‘children of the 90s’‐‐the index offspring of the Avon Longitudinal Study of Parents and Children. International Journal of Epidemiology. 2013; 42: 111–27.

17. Fraser A, Macdonald-Wallis C, Tilling K, Boyd A, Golding J, Davey Smith G, et al. Cohort Profile: The Avon Longitudinal Study of Parents and Children: ALSPAC mothers cohort. International Journal of Epidemiology. 2013; 42(1): 97–110.

18. Paternoster L HL, Tilling K, Weedon MN, Freathy RM, Frayling TM, Kemp JP, Davey Smith G, Timpson NJ, Ring SM, Evans DM, Lawlor DA. Adult height variants affect birth length and growth rate in children. Hum Mol Genet. 2011; 20: 4069–75.

19. Sieradzka D PR, Freeman D, Cardno AG, McGuire P, Plomin R, Meaburn EL, Dudbridge F, Ronald A. Are genetic risk factors for psychosis also associated with dimension-specific psychotic experiences in adolescence? PLoS ONE. 2014; 9: e94398.

20. Speliotes EK, Willer CJ, Berndt SI, Monda KL, Thorleifsson G, Jackson AU, et al. Association analyses of 249,796 individuals reveal 18 new loci associated with body mass index. Nat Genet. 2010;42(11):937–48.

21. Locke AE, Kahali B, Berndt SI, Justice AE, Pers TH, Day FR, et al. Genetic studies of body mass index yield new insights for obesity biology. Nature. 2015;518(7538):197–206.

22. O'Brien E, Asmar R, Beilin L, Imai Y, Mallion J-M, Mancia G, et al. European Society of Hypertension recommendations for conventional, ambulatory and home blood pressure measurement. Journal of Hypertension. 2003; 21(5): 821–48. P

23. Lang RM BM, Devereux RB, Flachskampf FA, Foster E, Pellikka PA, Picard MH, Roman MJ, Seward J, Shanewise JS, Solomon SD, Spencer KT, Sutton MS, Stewart WJ. Recommendations for chamber quantification: A report from the american society of echocardiography's guidelines and standards committee and the chamber quantification writing group, developed in conjunction with the european association of echocardiography, a branch of the european society of cardiology. J Am Soc Echocardiogr. 2005; 18: 1440–63.

24. Steeden JA, Atkinson D, Hansen MS, Taylor AM, Muthurangu V. Rapid Flow Assessment of Congenital Heart Disease with High-Spatiotemporal-Resolution Gated Spiral Phase-Contrast MR Imaging. Radiology. 2011; 260(1): 79–87.

25. Stergiopulos N MJ, Westerhof N. Simple and accurate way for estimating total and segmental arterial compliance: the pulse pressure method. Ann Biomed Eng. 1994; 22: 392–7.

26. OPCS. Standard Occupational Classification Volume 3. London: HMSO; 1991.

27. Anderson EL TK, Fraser A, Macdonald-Wallis C, Emmett P, Cribb V, Northstone K, Lawlor DA, Howe LD. Estimating trajectories of energy intake through childhood and adolescence using linear-spline multilevel models. Epidemiology. 2013; 24: 507–15.

28. Mattocks C NA, Leary S, Tilling K, Blair SN, Shield J, Deere K, Saunders J, Kirkby J, Smith GD, Wells J, Wareham N, Reilly J, Riddoch C. Use of accelerometers in a large field-based study of children: protocols, design issues, and effects on precision. J Phys Act Health. 2008;5:S98–S111.

29. Palmieri V, de Simone G, Arnett DK, Bella JN, Kitzman DW, Oberman A, et al. Relation of various degrees of body mass index in patients with systemic hypertension to left ventricular mass, cardiac output, and peripheral resistance (The Hypertension Genetic Epidemiology Network Study). The American Journal of Cardiology. 2001; 88(10): 1163–8.

30. Chinali M, Devereux RB, Howard BV, Roman MJ, Bella JN, Liu JE, et al. Comparison of cardiac structure and function in American Indians with and without the metabolic syndrome (the Strong Heart Study). The American Journal of Cardiology. 2004;93(1):40–4.

31. Palmer TM LD, Harbord RM, Sheehan NA, Tobias JH, Timpson NJ, Davey Smith G, Sterne JAC. Using multiple genetic variants as instrumental variables for modifiable risk factors. Stat Methods Med Res. 2012; 21: 223–42.

32. Richmond RC DSG, Ness AR, den Hoed M, McMahon G, Timpson NJ. Assessing causality in the association between child adiposity and physical activity levels: a Mendelian randomization analysis. PLOS Med. 2014;11:e1001618.

33. JH SDaS. Instrumental variables regression with weak instruments. Econometrica. 1997; 65: 557–86.

34. Baum C SM, Stillman S. IVENDOG: Stata module to calculate Durbin-Wu-Hausman endogeneity test after ivreg. In: S494401 SSC, editor. Boston: Boston Collge Department of Economics 2007.

35. Ferreira JP, Girerd N, Bozec E, Machu JL, Boivin JM, London GM, et al. Intima–Media Thickness Is Linearly and Continuously Associated With Systolic Blood Pressure in a Population-Based Cohort (STANISLAS Cohort Study). Journal of the American Heart Association. 2016;5(6).

36. Kim EJ, Park CG, Park JS, Suh SY, Choi CU, Kim JW, et al. Relationship between blood pressure parameters and pulse wave velocity in normotensive and hypertensive subjects: invasive study. J Hum Hypertens. 2006; 21(2): 141–8.

37. Malcolm DD, Burns TL, Mahoney LT, Lauer RM. Factors Affecting Left Ventricular Mass in Childhood: The Muscatine Study. Pediatrics. 1993; 92(5): 703–9.

38. Johnson W, Kuh D, Tikhonoff V, Charakida M, Woodside J, Whincup P, et al. Body Mass Index and Height From Infancy to Adulthood and Carotid Intima-Media Thickness at 60 to 64 Years in the 1946 British Birth Cohort Study. Arteriosclerosis, Thrombosis, and Vascular Biology. 2014; 34(3): 654–60.

39. de Simone G, Devereux RB, Daniels SR, Koren MJ, Meyer RA, Laragh JH. Effect of growth on variability of left ventricular mass: Assessment of allometric signals in adults and children and their capacity to predict cardiovascular risk. Journal of the American College of Cardiology. 1995; 25(5): 1056–62.

40. Stergiakouli E, Gaillard R, Tavaré JM, Balthasar N, Loos RJ, Taal HR, et al. Genome-wide association study of height-adjusted BMI in childhood identifies functional variant in ADCY3. Obesity (Silver Spring, Md). 2014; 22(10): 2252–9.

41. Bowden J, Davey Smith G, Burgess S. Mendelian randomization with invalid instruments: effect estimation and bias detection through Egger regression. International Journal of Epidemiology. 2015. 512–525.

42. Corbin LJ, Richmond RC, Wade KH, Burgess S, Bowden J, Smith GD, et al. Body mass index as a modifiable risk factor for type 2 diabetes: Refining and understanding causal estimates using Mendelian randomisation. Diabetes. 2016:65: 3002–7.

43. Bowden J, Davey Smith G, Haycock PC, Burgess S. Consistent Estimation in Mendelian Randomization with Some Invalid Instruments Using a Weighted Median Estimator. Genetic Epidemiology. 2016; 40(4): 304–14.

44. Hartwig FP, Davies NM. Why internal weights should be avoided (not only) in MR-Egger regression. International Journal of Epidemiology. 2016:45: 1676–1678.

45. Bowden FJ, Burgess S, Davey Smith G. Response to Hartwig and Davies. International Journal of Epidemiology. 2016:45: 1679–1680.

46. Yaghootkar H, Lotta LA, Tyrrell J, Smit RAJ, Jones SE, Donnelly L, et al. Genetic Evidence for a Link Between Favorable Adiposity and Lower Risk of Type 2 Diabetes, Hypertension, and Heart Disease. Diabetes. 2016; 65(8): 2448–60.

47. Charakida M JA, Falaschetti E, Khan T, Finer N, Sattar N, Hingorani A, Lawlor DA, Davey Smith G, Deanfield JE. Childhood obesity and vascular phenotypes: a population study. Journal of the American College of Cardiology. 2012; 60: 2643–50.

48. Lavie CJ AC, Ventura HO, Messerli FH. Left atrial abnormalities indicating diastolie ventricular dysfunction in cardiopathy of obesity. Chest. 1967; 92: 1042–6.

49. Vaziri SM LM, Lauer MS, Benjamin EJ, Levy D. Influence of blood pressure on left atrial size. The Framingham Heart Study. Hypertension. 1995; 25: 1155–60.

50. Burgess S, Timpson NJ, Ebrahim S, Davey Smith G. Mendelian randomization: where are we now and where are we going? International Journal of Epidemiology. 2015; 44(2): 379–88.

51. Gray L, Lee IM, Sesso HD, Batty GD. Blood pressure in early adulthood, hypertension in middle-age, and future cardiovascular disease mortality: the Harvard Alumni Health Study. Journal of the American College of Cardiology. 2011;58(23):2396–403.

52. Levy D, Garrison RJ, Savage DD, Kannel WB, Castelli WP. Prognostic Implications of Echocardiographically Determined Left Ventricular Mass in the Framingham Heart Study. New England Journal of Medicine. 1990; 322(22): 1561–6.

53. Dhuper S AR, Weichbrod L, Mahdi E, Cohen HW. Association of obesity and hypertension with left ventricular geometry and function in children and adolescents. Obesity. 2011; 19: 128–33.

54. Friedemann C, Heneghan C, Mahtani K, Thompson M, Perera R, Ward AM. Cardiovascular disease risk in healthy children and its association with body mass index: systematic review and meta-analysis. BMJ: British Medical Journal. 2012; 345: e4759.

55. Bogers RP, Bemelmans WE, Hoogenveen RT, et al. Association of overweight with increased risk of coronary heart disease partly independent of blood pressure and cholesterol levels: A meta-analysis of 21 cohort studies including more than 300 000 persons. Archives of Internal Medicine. 2007; 167(16): 1720–8.

56. Meyer AA, Kundt G, Steiner M, Schuff-Werner P, Kienast W. Impaired Flow-Mediated Vasodilation, Carotid Artery Intima-Media Thickening, and Elevated Endothelial Plasma Markers in Obese Children: The Impact of Cardiovascular Risk Factors. Pediatrics. 2006; 117(5): 1560–7.

57. Woo KS, Chook P, Yu CW, Sung RYT, Qiao M, Leung SSF, et al. Overweight in children is associated with arterial endothelial dysfunction and intima-media thickening. Int J Obes Relat Metab Disord. 2004; 28(7): 852–7.

58. Iannuzzi A, Licenziati MR, Acampora C, Salvatore V, Auriemma L, Romano ML, et al. Increased Carotid Intima-Media Thickness and Stiffness in Obese Children. Diabetes Care. 2004; 27(10): 2506.

59. Urbina EM, Kimball TR, Khoury PR, Daniels SR, Dolan LM. Increased Arterial Stiffness is Found in Adolescents with Obesity or Obesity-Related Type 2 Diabetes Mellitus. Journal of hypertension. 2010; 28(8): 1692–8.

60. Corden B, Keenan NG, de Marvao ASM, Dawes TJW, DeCesare A, Diamond T, et al. Body Fat Is Associated With Reduced Aortic Stiffness Until Middle Age. Hypertension. 2013; 61(6): 1322–7.

61. Sorof JM, Poffenbarger T, Franco K, Bernard L, Portman RJ. Isolated systolic hypertension, obesity, and hyperkinetic hemodynamic states in children. The Journal of Pediatrics. 2002; 140(6): 660–6.

62. Jiang X, Srinivasan SR, Urbina E, Berenson GS. Hyperdynamic Circulation and Cardiovascular Risk in Children and Adolescents. Circulation. 1995; 91(4): 1101–6.

63. Smith GD, Lawlor DA, Harbord R, Timpson N, Day I, Ebrahim S. Clustered Environments and Randomized Genes: A Fundamental Distinction between Conventional and Genetic Epidemiology. PLOS Medicine. 2007; 4(12): e352.

64. Cole SR, Platt RW, Schisterman EF, Chu H, Westreich D, Richardson D, et al. Illustrating bias due to conditioning on a collider. International Journal of Epidemiology. 2010; 39(2): 417–20.

65. Aschard H, Vilhjálmsson Bjarni J, Joshi Amit D, Price Alkes L, Kraft P. Adjusting for Heritable Covariates Can Bias Effect Estimates in Genome-Wide Association Studies. American Journal of Human Genetics. 2015; 96(2): 329–39.

